# Archaeal G-Quadruplexes: A Novel Model for Understanding Unusual DNA/RNA Structures Across the Tree of Life

**DOI:** 10.1101/2024.01.16.575881

**Authors:** Zackie Aktary, Kate Sorg, Anne Cucchiarini, Guglielmo Vesco, Dorian Noury, Rongxin Zhang, Thomas Jourdain, Daniela Verga, Pierre Mahou, Nicolas Olivier, Natalia Bohalová, Otilia Porubiaková, Václav Brázda, Marie Bouvier, Marta Kwapisz, Béatrice Clouet-d’Orval, Thorsten Allers, Roxane Lestini, Jean-Louis Mergny, Lionel Guittat

## Abstract

Archaea, a domain of microorganisms found in diverse environments including the human microbiome, represent the closest known prokaryotic relatives of eukaryotes. This phylogenetic proximity positions them as a relevant model for investigating the evolutionary origins of nucleic acid secondary structures such as G-quadruplexes (G4s), which play regulatory roles in transcription and replication. Although G4s have been extensively studied in eukaryotes, their presence and function in archaea remain poorly characterized. In this study, a genome-wide analysis of the halophilic archaeon Haloferax volcanii identified over 5, 800 potential G4-forming sequences. Biophysical validation confirmed that many of these sequences adopt stable G4 conformations in vitro. Using G4-specific detection tools and super-resolution microscopy, G4 structures were visualized in vivo in both DNA and RNA across multiple growth phases. Comparable findings were observed in the thermophilic archaeon Thermococcus barophilus. Functional analysis using helicase-deficient H. volcanii strains further identified candidate enzymes involved in G4 resolution. These results establish H. volcanii as a tractable archaeal model for G4 biology.

## INTRODUCTION

Archaea, one of the three domains of life, are distinct from bacteria and eukaryotes and represent a diverse and fascinating group of microorganisms that often thrive in extreme environments (1, 2). Unlike bacteria and eukaryotes, archaea possess unique cell membrane lipids—typically ether-linked isoprenoids—and lack peptidoglycan in their cell walls, a feature common in bacteria (3). Archaea provide valuable insights into the evolutionary history of life on Earth. Notably, the discovery of Asgard archaea supports the emerging “two-domain” tree of life, suggesting that eukaryotes evolved from the archaeal lineage (4–6).

Archaea’s unique characteristics make them a subject of ongoing interest across fields like microbiology, biotechnology, and astrobiology. Many archaea are extremophiles, inhabiting environments hostile to most other life forms, such as high-temperature (thermophiles), high acidity or alkalinity (acidophiles and alkaliphiles), high-salinity (halophiles), or high-pressure (piezophiles) ecosystems (3). Beyond extreme habitats, archaea also play a significant, yet less understood, role in the human microbiome, particularly in the gastrointestinal tract, skin, and oral cavity (7, 8). They are central to various biogeochemical cycles, such as the carbon and nitrogen cycles, and are key players in methanogenesis, an anaerobic process contributing to both natural and human-related methane emissions. Extremophilic archaea are also important in biotechnology where their enzymes (*e.g.*, DNA polymerases, lipases, and esterases) have industrial applications (9).

Although distinct from eukaryotes, archaea share more molecular and genetic similarities with eukaryotic organisms than with bacteria, including the presence of histones or histone-like proteins, chromatin-like structures, and complex DNA/RNA polymerases, underscoring their evolutionary significance (10, 11). The study of archaea provides critical insights into fundamental cellular processes such as replication, transcription, translation, and DNA repair, shedding light on our evolutionary past (4, 12). In addition to the canonical double-helix structure, DNA can adopt non-canonical configurations, at least transiently (13–15). Among these, G-quadruplexes (G4s) are unique four-stranded nucleic acid structures found both at the DNA and RNA levels (16, 17). G4s are formed by the stacking of four guanine (G) bases into square planar arrangements called G-quartets. Over the past few decades, G4s have garnered significant attention for their roles in cellular processes and potential as therapeutic targets (18–20). These structures adopt various topologies depending on factors like sequence, loop length, and ionic conditions. Monocations, such as potassium (K^+^) or sodium (Na^+^), influence their structure, stability, and biological functions (21–23). G4s are implicated in critical biological processes, including transcription, telomere maintenance (24), DNA replication (25, 26), DNA repair (27, 28), RNA processing, and translation (29, 30). As regulatory elements, G4s act as molecular switches, modulating gene expression and protein-DNA interactions (31–33). Recent evidence suggests G4s may also contribute to epigenetic regulation, influencing cell-type-specific transcriptomes (34, 35) and higher-order chromatin structures (31, 36). These unusual structures have emerged as promising therapeutic targets for diseases such as cancer, viral infections, and neurodegenerative disorders (20, 37, 38). Numerous G4-stabilizing ligands (G4 ligands or G4L) have been identified, and some are under investigation for their therapeutic potential (18, 39–41).

The evolutionary history of G4s is still poorly understood and investigated (42). G4s have been extensively studied in various organisms from viruses(43–45), bacterias(46, 47) to eukaryotes with the special effort for analyses of the human genome(48–50). While G4s in bacteria have been linked to radioresistance, antigenic variation, recombination, and the regulation of gene expression, their presence and biological relevance in archaea remain largely uncharacterized (47, 51). A promising tool to study and characterize G4s in archaea is super-resolution microscopy, which has already been used to visualize nascent DNA replication foci in archaeal cells (52, 53).

A significant limitation of conventional optical microscopy is its inability to distinguish simultaneous emissions from two adjacent fluorophores within a diffraction-limited area. This restricts its resolution to molecular features larger than 250 nm. Over the past two decades, super-resolution microscopy techniques have been developed to address this limitation. These methods, such as Structured Illumination Microscopy (SIM, resolution 100-150 nm) (54) and Single-Molecule Localization Microscopy (SMLM, resolution 10-40 nm) such as Stochastic Optical Reconstruction Microscopy (STORM) (55), offer greatly improved resolution (56). Technical advances in super-resolution microscopy have been instrumental in advancing our understanding of processes like DNA replication, transcription, and damage response pathways, which are challenging to study using conventional microscopy or ensemble-based assays.

In this study, we investigated the existence and potential implications of G4 structures in archaeal genomes, focusing on *Haloferax volcanii*, a salt-loving archaeal model (57). Using the G4Hunter prediction tool, we analyzed differences in the presence and frequency of potential G4-forming sequences (PQS) in the *H. volcanii* genome (58). Our bioinformatics predictions were experimentally validated *in vitro* using an array of biophysical methods and *in vivo* in archaeal cells using the G4-specific antibody BG4 (59). Using different super-resolution microscopy techniques, we demonstrate, for the first time, that G4s are present in both the DNA and RNA in archaea. Their presence can be dynamically modulated by G4-specific ligands, which stabilize these structures, and by helicases, which actively unwind G-quadruplexes.

## MATERIAL AND METHODS

### Analytical process

All sequences were analyzed using the G4Hunter Web tool (http://bioinformatics.ibp.cz/#/analyse/quadruplex), after extracting National Center for Biotechnology Information (NCBI) IDs. Unless specified otherwise, parameters for G4Hunter were set to 25 nucleotides for window size and a threshold score of at least 1.2. This threshold appears as a reasonable compromise, giving few false positives (sequences not forming a G4 despite a G4Hunter score above threshold) and false negatives (sequences able to form a stable G4 despite having a G4Hunter score below threshold). Scores above 1.2 correspond to sequences having a relatively high guanine content and likely to form stable G4s. To rank sequences based on score, motifs were binned in five intervals covering the G4Hunter scores 1.2–1.4, 1.4–1.6, 1.6–1.8, 1.8–2.0 and >2.0 (58, 103). The higher the score, the higher the probability to form a stable G4.

### G4 prediction in *Haloferax volcanii* promoters

The G4Hunter software was used to predict all potential G4s in the promoter regions of *Haloferax volcanii*. The window size was set to 25 bp, and the score threshold was set to 1.5. The promoter region was defined as the 70 bp region upstream of the transcription start site (TSS) (62), and G4Hunter was used to scan this region. The EnrichedHeatmap package (V 1.32.0) was utilized to calculate the density of G4s within 500 bp upstream and downstream of the TSS, employing the coverage mode with a calculation resolution of 1 bp.

### Oligonucleotide sequences

Oligonucleotides were purchased from Eurogentec (Belgium) and used without further purification. F21T, the fluorescently labelled reference quadruplex, negative and positive controls, as well as the tested sequences were prepared at 100 μM strand concentration in ddH2O. Sequence details are provided in Supplementary Tables 1, 2 and 3. All oligonucleotides were annealed (95°C for 5 min and slowly cooled to room temperature) in the corresponding buffer before measurements.

### Circular dichroism

3 μM oligonucleotide solutions were annealed in K100 buffer (10 mM lithium cacodylate, 100 mM KCl, pH 7.2). CD spectra were recorded on a J-1500 spectropolarimeter (JASCO, France) at room temperature (25°C), using a scan range of 350–210 nm, a scan rate of 100 nm/min and averaging 3 accumulations.

### FRET-melting competition assay

FRET melting competition (FRET-MC) experiments were performed in 96-well plates using a CFX96 qPCR Real-Time PCR (Biorad). The tested sequences (or competitors) were annealed at 7.5 µM in K10 buffer (10 mM KCl, 10 mM lithium cacodylate, 90 mM LiCl, pH 7.2) and F21T was annealed at 5 µM in K10. Each well contained 3 µM competitors, 0.2 μM fluorescent oligonucleotide F21T with or without 0.4 μM G4 ligand (PhenDC3) in K10 buffer, for a total volume of 25 μL. Samples were kept at 25°C for 5 min, then the temperature was increased by 0.5°C per minute until 95°C, and the FAM channel was used to collect the fluorescence signal. The melting temperature (Tm) of an oligonucleotide is defined as the temperature where 50% of the oligonucleotide is unfolded and corresponds to the normalized value of 0.5. ΔTm is determined by subtracting the Tm containing PhenDC3 to the Tm in absence of PhenDC3. The S-Factor, an indicator of G4 formation, was calculated as previously described (65). The FRET-MC assay uses a 96-well plate as a sample holder, which allows to process 48 sequences simultaneously. Each experimental condition was tested in duplicate on two separate plates.

### Isothermal difference spectra (IDS)

Absorbance spectra were recorded on a Cary 300 spectrophotometer (Agilent Technologies, France) (scan range: 330-220 nm; scan rate: 600 nm/min; automatic baseline correction). 3 μM oligonucleotide solutions were annealed in Li-CaCo10 buffer (10 mM lithium cacodylate, pH 7.2). Absorbance spectra were first recorded at 25°C in the absence of any stabilizing cation. KCl was added after recording the first spectrum, to reach a final potassium concentration of 100 mM. The second UV-absorbance spectrum was recorded after 15 min of equilibration. IDS corresponds to the arithmetic difference between the initial (unfolded) and second (folded, thanks to the addition of K+) spectra, after correction for dilution.

### G-Quadruplex fluorescent light-up probes

Thioflavin T (ThT) was used as previously described (66). The tested oligonucleotides were annealed at 7.5 µM in K100 buffer. The oligonucleotides and ThT (dissolved in milli Q water) were added in a microplate at 3 and 2 µM respectively, to reach a final volume of 100 µL in K100 buffer. The plate was shaken for 10 sec and was incubated for 10 min at room temperature. Fluorescence intensity was collected at 490 nm after excitation at 420 nm in a TECAN M1000 pro plate reader. The fluorescence signal was normalized to the oligonucleotide length. Each condition was done in duplicate on different plates. NMM (N-methyl mesoporphyrin IX) was used under the same conditions as ThT, except that fluorescence intensity was collected at 610 nm after excitation at 380 nm in a TECAN M1000 pro plate reader.

### Strains, plasmids and growth conditions

H. volcanii cultures using enriched Hv-YPC media were grown at 45°C, as described previously (104). Hv-YPC contained (per liter) 144 g of NaCl, 21 g of MgSO4.7H2O, 18 g of MgCl2.6H2O, 4.2 g of KCl, and 12 mM Tris–HCl (pH 7.5), Yeast extract (0.5%, wt/vol; Difco), 0.1% (wt/vol) peptone (Oxoid), and 0.1% (wt/vol) Casamino Acids (Difco). For experiments using the G4L PhenDC3, overnight cultures were diluted to OD600nm of 0.1 and incubated at 45°C. PhenDC3 (Sigma) was added (to a final concentration of 10 µM) after one hour of growth, and the treatment was continued for two hours at 45°C. An OD600nm between 0.1 and 0.2 was used for exponential phase cells, whereas OD600nm of 1.2 was used for stationary phase.

Cells of Thermococcus barophilus (TbΔ517; UBOCC-M-3300;(105)) were grown at 85°C under atmospheric pressure in anaerobic conditions (50 mL penicillin vials) in Thermococcales rich medium (TRM; (106)) supplemented with elemental sulphur (0.25 g/L; (70)). To determine growth, cells were counted using a Thoma counting chamber (0.01 mm depth, Weber, England). For exponential phase, cells were grown for 6h after inoculation from saturated pre-culture (1×10^7^ cells/mL) and for stationary phase for 18h (1×10^8^ cells/mL).

### Construction of strains with deletions of Rad3 and Dna2 helicases

To generate the Δdna2 construct (strain H431), the upstream regions (US) of dna2 and the down-stream regions (DS) of dna2 were cloned into pTA131 to generate the pTA417 plasmid, which was used to construct the deletion strain by a gene knockout system (44). The dna2 deletion was generated in the H53 strain (ΔpyrE2 ΔtrpA) (104). The US and DS regions were generated by PCR on H26 genomic DNA (ΔpyrE2) using specific primers. The template DNA used was isolated from genomic DNA libraries of H. volcanii using internal primers with BamHI sites and external primers with XbaI and KpnI sites, inserted at KpnI and XbaI sites in pTA131 plasmid.

To generate the Δrad3a and Δrad3b constructs (strains HvLG1 and HvLG2, respectively), the upstream regions (US) of rad3a/b and the down-stream regions (DS) of rad3a/b were cloned into pTA131 to generate the pLG1/2 plasmids, which were used to construct the deletion strains by a gene knockout system (107). The rad3a/b deletion was generated in the H26 strain (ΔpyrE2) (104). The US and DS regions were generated by PCR on H26 genomic DNA using specific primers. Each PCR product contained 30 bp homology with adjacent fragments for sequence and ligation independent cloning (SLIC) (108). Following the SLIC method, PCR fragments and linearized plasmid were digested by T4 DNA polymerase exonuclease activity for 45 min at 22°C to generate 3’-single-stranded extremities, then all amplification products were mixed in a 1:1:1 molecular ratio and incubated 30 min at 37°C prior to transformation into E. coli XL1-blue cells. The presence of the correct inserts was tested by PCR amplification on white colonies using specific primers. The sequence of selected plasmids, pLG1 and 2, were further confirmed by sequencing on both strands. Both plasmids were used to transform strain H26 (ΔpyrE2) using the pop-in/pop-out method as described previously (107). The absence of the rad3a and rad3b genes was tested by colony lift on 100 colonies from the pop-out plates using a probe targeting rad3a or rad3b. The digoxigenin (DIG)-labelled probes were generated by PCR on H26 genomic DNA using primers and the PCR DIG labelling Mix from Roche. Probe hybridization was detected using the DIG Luminescent Detection kit (Roche) and a ChemiDoc MP (BioRad). All the oligonucleotides used for the different constructions are listed in Supplemental Table 3. The double Δrad3a Δrad3b mutant (strain HvLG3) was constructed by deleting rad3b in the Δrad3a mutant.

### In vivo visualization of G4 structures: 3D-Structured Illumination microscopy (3D-SIM) and STORM

*H. volcanii* cells were fixed with 2 % formaldehyde (FA) for 30 min at 45°C under agitation followed by the addition of glycine (0.5M) for 20 minutes at 45°C under agitation. Cells were washed with PBS, permeabilized with 60% ethanol, washed and resuspended in 1 ml PBS for cultures with an OD600nm of 0.2. 20 µl of cell solution was spotted on poly-L-Lysine coated coverslips for 30 minutes, washed with PBS and incubated with 100 µg/ml RNaseA (ThermoFisher) for 30 min at 37°C. Coverslips were then washed with PBS and blocked with blocking solution (PBS + 5% goat serum + 0.1% Tween) for 30-60 minat room temperature. Next, coverslips were incubated with the BG4 antibody (10 nM) for 1 h. Coverslips were then washed with PBS + 0.1% Tween (PBS-Tween) and incubated with a mouse anti-FLAG antibody (Sigma ref) (1/1000) for 45 min. After washing with PBS-Tween, slides were incubated with anti-Mouse IgG-DyLight (ref) (1/1000) for 45 min (DyLight 549 for SIM and DyLight755 for STORM). All incubations were done at room temperature and all antibodies were prepared in antibody solution (PBS + 1% goat serum + 0.1% Tween). Coverslips were washed twice with PBS-Tween, then incubated with Yoyo1 (Invitrogen, 1/10 000) for 10 minutes at room temperature and washed two times with PBS-Tween followed by two washes with PBS. Images were acquired on a DeltaVision OMX SR imaging system from GE Healthcare (Velizy, France) with an Olympus PlanApo N 60× 1.42 NA oil immersion objective. The quality of the raw SI data was checked by the SIMcheck suite of plugins available for Fiji software prior to SI reconstruction performed with the softWoRx software from Applied Precision. Image analysis was performed using custom scripts in Napari (109). Briefly, individual cell outlines were segmented using the nucleic acid signal and Cellpose pretrained models (110). In each cell, the integrated signal and other features were determined using the scikit-image package (111).

STORM imaging was performed on an IX83 Inverted microscope (Olympus) using a 100x 1.3NA objective (Olympus) and an Orca Fusion sCMOS camera (Hamamatsu) using the slowest read-out speed and 2x binning, resulting in a pixel size of 130nm. We used a 639 nm laser (Voltran, 100mW) for excitation, at a power of ∼1kW/cm2. The microscope is equipped with Chroma filters: the fluorophores were imaged using a ZT532/640rpc 2-color dichroic mirror and a ET700-75 emission filter. Due to the double-deck design, the emitted light also goes through a T660lpxr dichroic mirror. The sample were imaged in an Attofluor imaging chamber (Invitrogen, A7816), with ∼1ml of imaging buffer and another 25mm round coverglass on top to limit air exchanges. The STORM buffer was prepared according to (112).

Raw STORM image stacks were processed using a FIJI macro that runs an analysis with Detection of Molecules (DoM) (113) including drift-correction and grouping of consecutive localizations. (available at https://github.com/LaboratoryOpticsBiosciences/STORM_Analysis)

#### BG4 purification

The BG4-encoding plasmid (Plasmid #55756, Addgene), was transformed into Escherichia coli strain Rosetta 2 pLys (Novagen), followed by culturing in 2xYT and kanamycin (50 μg/ml) and chloramphenicol (34 μg/ml) at 37°C and 250 RPM. When OD600nm reached 0.6–0.8 the temperature was lowered to 20°C, and induction was initiated with 0.5 mM IPTG. After 15 h of induction at 20°C, the bacterial culture was harvested by spinning at 4, 000 g for 30 min. The resulting pellet was resuspended in 8 ml TES buffer 1 (50 mM Tris-HCl pH 8.0, 1 mM EDTA pH 8.0, 20% sucrose) per 100 ml volume expression culture. The mixture was put on ice for 10 min. Next, 12 ml of TES buffer 2 (10 mM Tris-HCl pH 8.0, 0.2 mM EDTA pH 8.0, 4% sucrose, with benzonase nuclease and 2 mM MgSO4) per 100 ml volume expression culture was added and the mixture was put on ice for 15 min. Cells were centrifuged at 4°C for 20 min at 4000 g. The supernatant was collected and filtered (0.22 μm). The sample was deposited on a washed and PBS-preequilibrated HisTrap HP column (Cytiva). The resin was then washed with 20 column volumes (CV) of high salt wash buffer (PBS, 350 mM NaCl, 10 mM imidazole pH 8.0) and 20 CV low salt wash buffer (PBS, 10 mM imidazole pH 8.0). The BG4 protein was eluted with elution buffer (PBS, 500 mM imidazole pH 8.0), buffer-exchanged to PBS (Econo-Pac 10DG Column, Biorad) and concentrated using 50 ml centrifugal devices with 10 kDa cut-off (Millipore). Protein quality was assessed by running SDS-PAGE and visualized with Coomassie staining. Final protein concentration was calculated based on absorbance measurements at 280 nm.

## RESULTS

### Potential quadruplex sequences in *Haloferax volcanii* genomic DNA

To assess the potential for G4 formation in the *H. volcanii* genome, we first used the G4Hunter algorithm to determine the presence of potential quadruplex sequences (PQS) both in chromosomal DNA (chr) and in the four plasmids (pHV1-4). Using standard values for G4Hunter (i.e., a window size of 25 nucleotides and threshold score of 1.2), we found 5812 PQS in the entire genome of *H. volcanii*. The GC content is above 50% for all *H. volcanii* genomes, with a maximum for the main chromosome and pHV3 (around 66%), followed by pHV4 (61%), pHV2 (56%), and pHV1 (55%). Total PQS counts, percentage of GC and PQS frequencies are summarized in Figure 1. The vast majority of these PQS are present in the chromosome (4406), then in the two mini-chromosomes (723 and 605, respectively, for pHV4 and pHV3). The pHV1 contains 68 PQS and the smallest plasmid, pHV2, only 10 PQS. The highest PQS frequencies were found in the main chromosome and pHV2 (PQS around 1.55 per kb) followed by pHV3 and pHV4 (PQS 1.3 and 1.1, respectively). In contrast, the lowest PQS frequency was found in pHV1 (PQS = 0.8). Our bioinformatics analysis was performed on the wild-type strain, but it is worth noting that the plasmid pHV2 has been cured from the laboratory strain (H26). Moreover, in this strain, pHV4 is inserted in the main chromosome, leading to an additional and functional origin of replication (57, 60).

**Fig. 1.**
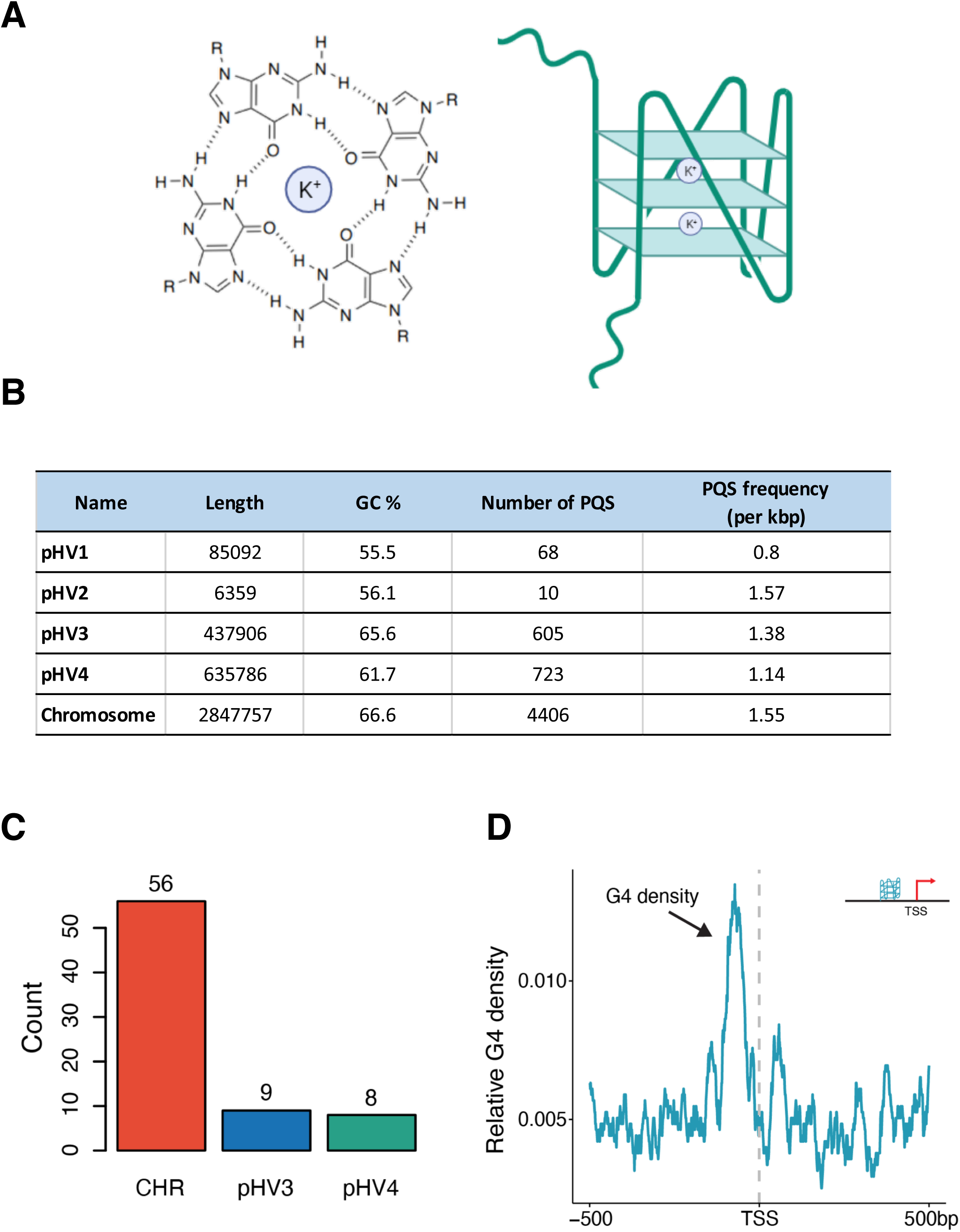
GC content, total number and their frequencies of PQS in *H. volcanii* genome. (A) Schematic presentation of a G-quartet (left) and a G-quadruplex (right). (B) Total PQS counts, percentage of GC and PQS frequency characteristics for the main chromosome and mini-chromosomes. (C, D) G4 prediction in *H. volcanii’s* promoters: number and localisation relative to the TSS.

The vast majority (98.5%) of PQS in the *H. volcanii* genome had a G4Hunter score between 1.2 and 1.4, which is relatively low (58). However, this finding is consistent with all genomes tested to date, including viruses, bacteria, archaea and eukaryotes (61). In *H. volcanii*, we investigated whether there was a correlation between the GC content of each genomic element (chromosome + mini-chromosomes) and the PQS frequency. We excluded pHV2 from our analysis because its size is very small (6kb) compared to the other components of the genome. We observed a clear correlation between GC content and PQS frequency between the 4 genomic elements (Chr, pHV1, pHV3, pHV4) (Supplementary Material 1).

The presence of G4 in promoters was found to be crucial in higher eukaryotes as they can regulate gene expression by influencing DNA stability, transcription factor binding, and the recruitment of the transcriptional machinery. We thus investigated the distribution and characteristics of PQS upstream of transcriptional start sites (TSSs) in *H. volcanii* that were previously identified by differential RNA-Seq (dRNA-Seq) (62). The chromosome contains a markedly higher number of potential promoter G4s (n = 56), compared to plasmids pHV3 (n = 9) and pHV4 (n = 8). This difference is likely attributed to the larger size and complexity of the chromosome relative to the plasmids.

We further examined the relative density of G4s near transcription start sites (TSSs), highlighting their spatial distribution. A prominent enrichment of G4s is observed upstream of the TSS, peaking sharply at approximately -100 bp. This enrichment underscores the potential regulatory significance of G4s in promoter regions in manner similar to mammalian cells (63).

### Experimental evidence for G4 formation *in vitro*

The G4Hunter false positive rate for sequences with scores above 1.5 is extremely low (nearly all candidate sequences form stable G4s *in vitro*). However, false positives may be observed for sequences with slightly less favorable scores, in the 1.2 to 1.5 interval (58). Since we observed that the majority of G4Hunter scores in *H. volcanii* were between 1.2 and 1.4, we aimed to verify, experimentally, that some of these identified sequences can adopt the G4 structure *in vitro*. To this end, we selected a panel of 36 sequences, containing four consecutive blocks of at least two guanines, distributed both on the chromosome and on the mini-chromosomes. To demonstrate the formation of the G4s, we used a combination of methods (64). FRET-MC enables the testing of several sequences in a parallel fashion (65). This approach is used to measure the ability of a test sequence to compete with the G4 ligand PhenDC3, which is highly G4-selective, for binding to a fluorescent G4-containing probe, F21T. Binding of PhenDC3 to the F21T stabilizes its G4, and increases the T_m_ of F21T. If the test sequence is forming a G4, it will act as “decoy” and trap PhenDC3, which will be less available to stabilize F21T. This competition will result in a less prominent stabilization of the T_m_ of F21T (ΔT_m_). Positive and negative controls (sequences known to form or not form G4s) were used for comparison. As shown in Figure 2A, 25 of the 36 sequences behaved as efficient competitors, suggesting that the majority of the motifs tested form G4s *in vitro*.

**Fig. 2.**
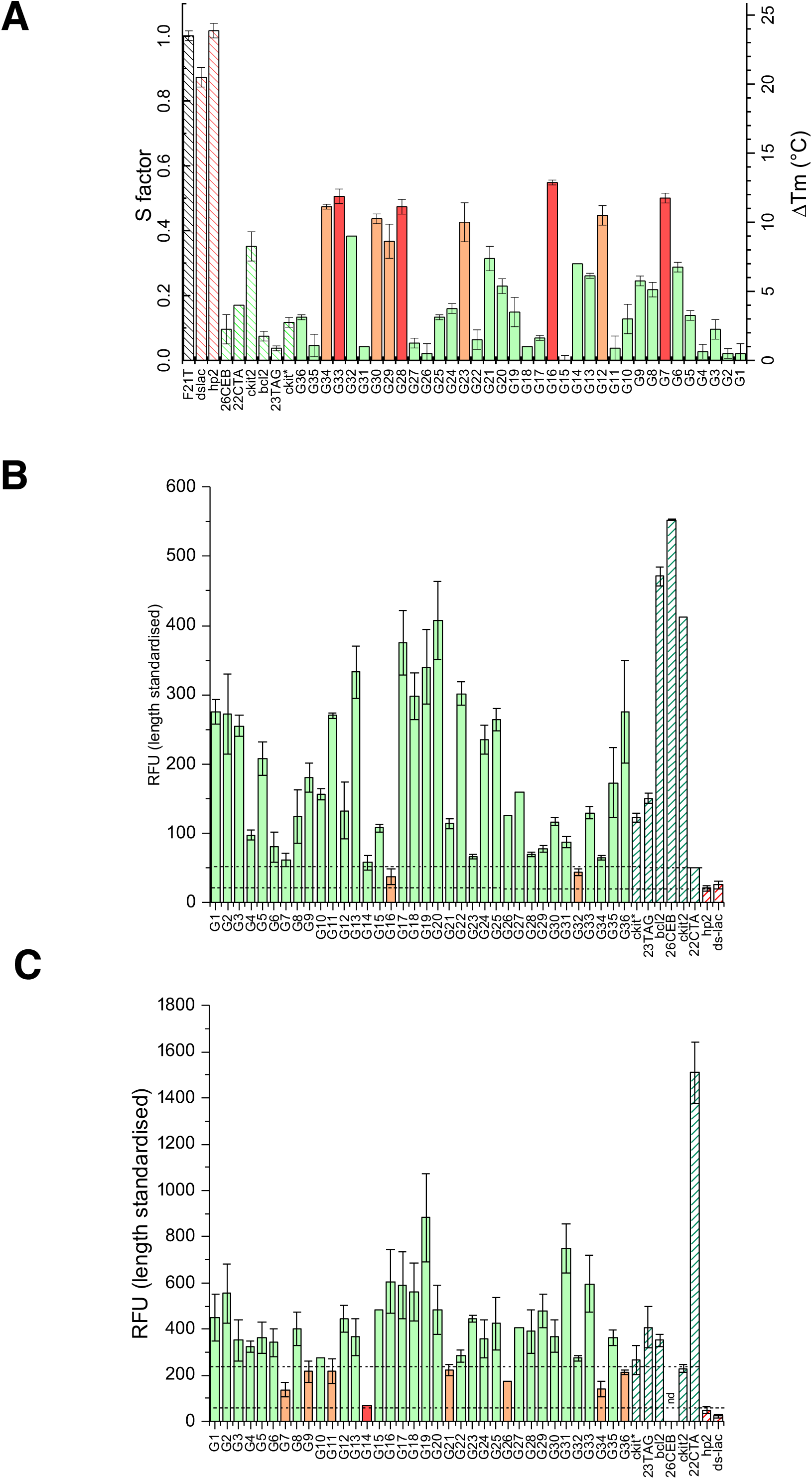
Evidence for G4 formation in 36 *H. volcanii* sequences. (A) FRET MC results. The S-factor is a score reflecting G4 formation. The lower the S-factor, the higher the G4 formation. (B) Thioflavin T fluorescence emission and (C) NMM fluorescence emission. Both Thioflavin T and NMM are light-up probes for which fluorescence intensity increases in the presence of G4-forming sequences. The color of the bars reflects the possibility of the sequences to be structured in G4 (green = yes, orange =likely, red = no).

As several independent techniques are recommended to definitively demonstrate G4 formation (64), we used an alternative approach and investigated whether the fluorescence emission of G4 probes such as thioflavin T (66) and NMM (67) was increased in the presence of potential G4-forming *H. volcanii* sequences. When compared to the negative and positive controls, most sequences induced an increase in fluorescence emission of both G4 light-up probes (Figure 2B and 2C). Finally, we performed additional spectroscopic experiments: isothermal difference spectra (IDS) and circular dichroism (CD). IDS experiments allow to compare the absorbance characteristics of the same oligonucleotide in its folded and unfolded forms. The arithmetic difference between the two corresponding spectra results in the isothermal difference spectra. Unfolding can be obtained by excluding stabilizing cations. G4s display a negative peak around 295 nm and a positive peak around 273 nm (Supplementary Figure S1). CD is widely used to characterize G-quadruplex (G4) structures by providing information on their folding topology—such as parallel, antiparallel, or hybrid conformations— based on their distinct CD spectral signatures. Most spectra obtained by CD were indicative of parallel G4 formation (Supplementary Figure S2). Altogether, these experiments confirmed that at least 26 of the 36 *H. volcanii* sequences tested fold into G4 conformations *in vitro* (Supplementary Table 4).

### G4s are present in archaea *in vivo*

We next aimed to validate our bioinformatic and *in vitro* results by demonstrating that G4s were present in archaeal cells *in vivo*. Using 3D-SIM super-resolution imaging, we performed immunofluorescence experiments using the G4-specific single-chain antibody (scFv) BG4 on fixed *H. volcanii* cells (59, 68). Given the density of G4 foci, we needed a higher resolution than the one provided by conventional fluorescence microscopy techniques (250 nm). 3D-SIM microscopy offers a resolution in the order of 100 nm (56). BG4 staining of exponentially growing *H. volcanii* cells revealed the presence of distinct, punctate foci distributed evenly throughout the cells while no staining was detected in the absence of BG4 (Figure 3A and B). These foci were present in *H. volcanii* cells in both exponential and stationary phases of growth, albeit to a significantly higher degree in exponential phase, suggesting the facilitated formation of G4 during genomic DNA replication. Therefore, we used exponential phase cells for our subsequent experiments (Figure 3C).

**Fig. 3.**
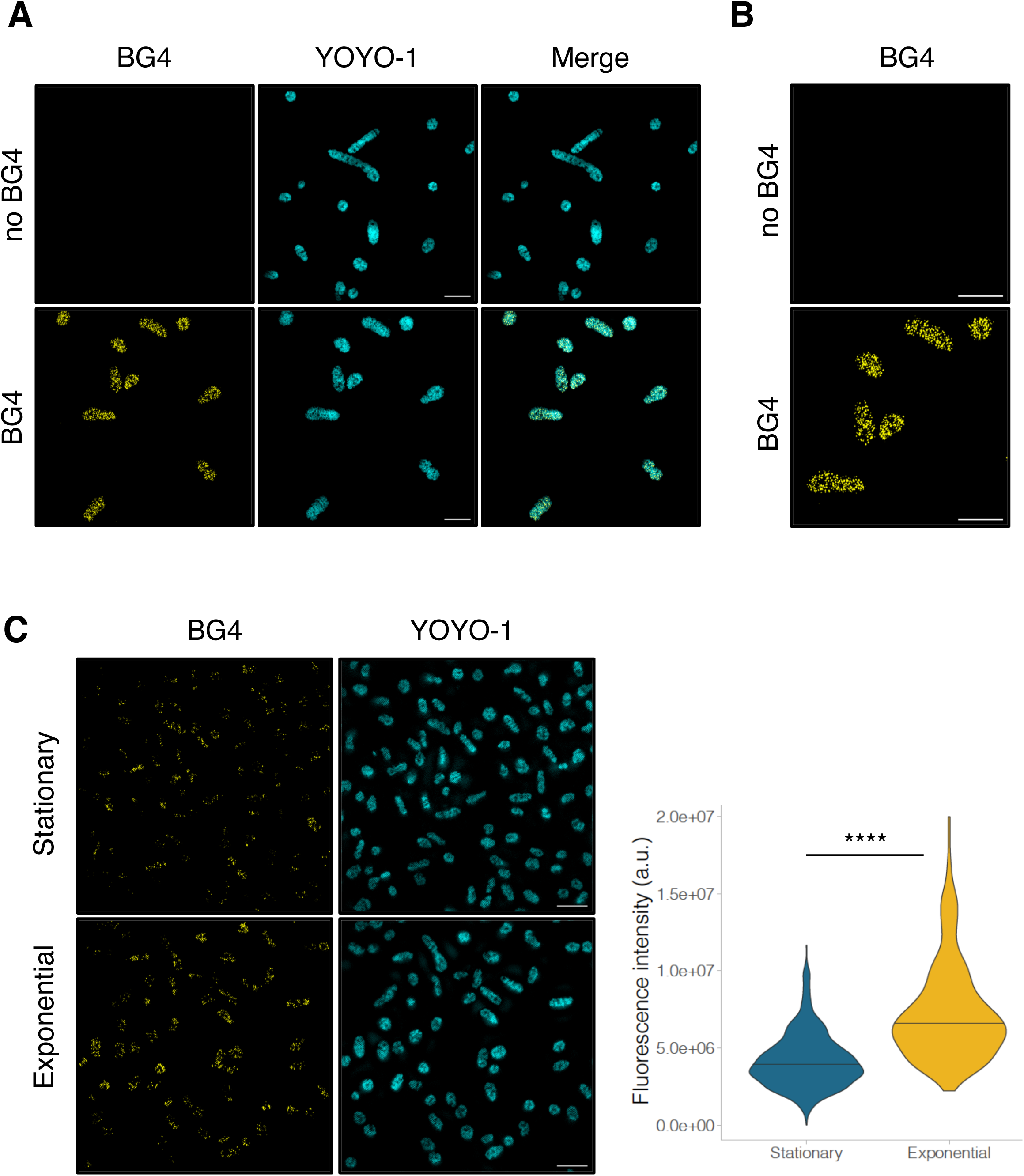
Immunofluorescence detection of G4s in *H. volcanii* by SIM. (A) Immunofluorescence labelling of G4 in *H. volcanii* using the BG4 antibody. YOYO-1 was used to visualize nucleic acids. (B) Magnification of BG4-stained *H. volcanii* cells. (C) Exponential and stationary growth phase cells. The number of samples per condition is: Exponential=273, Stationary=619. The summary of the data is shown as a violin plot reflecting the data distribution and a horizontal line indicating the median Bar, 5 µm.

Using transcriptomic and *in vitro* analyses, Guo and Bartel showed that G4 in RNA (rG4s) are depleted in the *E. coli* genome, most likely due to the lack of an efficient machinery to unwind them (69). To determine whether the G4s present in *H. volcanii* were being formed at the RNA or DNA level or both, we treated the cells with RNase A before staining with the BG4 antibody. While we were able to detect G4 structures in *H. volcanii* cells after RNase treatment, the G4 signal decreased approximately 1.7-fold compared to untreated cells (Figure 4A) demonstrating that a significant fraction (but not all) of the staining resulted from RNA G4 recognition.

**Fig. 4.**
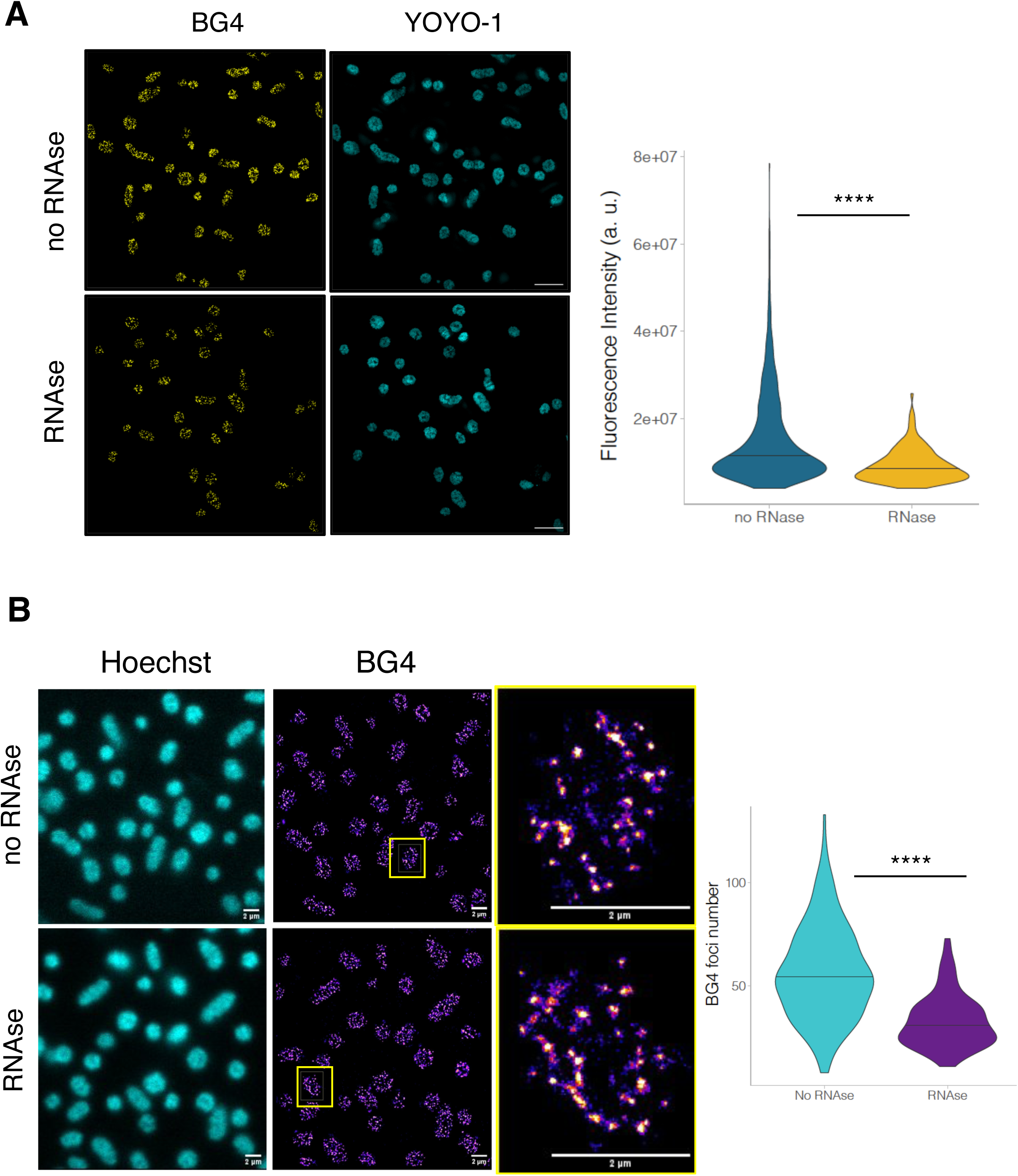
Immunofluorescence labelling and quantification of total G4 fluorescence intensity and G4 foci in *H. volcanii* by SIM and STORM. (A) Immunofluorescence labelling of G4 in *H. volcanii* using the BG4 antibody with or without RNase treatment by SIM. The number of samples per condition is: RNase=1120, no RNase=1095. YOYO-1 was used to visualize nucleic acids. (B) Immunofluorescence labelling of G4 in *H. volcanii* using the BG4 antibody with or without RNase treatment by STORM. Hoechst was used to visualize DNA. The number of samples per condition is: No RNase=169, RNase=149. Statistical analyses were performed using a Mann-Whitney test. **** p > 0.0001.

SIM was primarily employed in our study as it is a remarkable technique, offering excellent resolution and versatility for imaging BG4 foci. However, for specific cases requiring even higher spatial resolution, we complemented our analysis with STORM to ensure more precise quantification of BG4 foci, thereby leveraging the strengths of both approaches.

The detection of BG4 foci in *H. volcanii* cells, both with and without RNase treatment, confirmed evidence for the presence of G-quadruplex structures in both DNA and RNA (Figure 4B). Under RNase-free conditions, a higher number of BG4 foci were observed, suggesting the coexistence of G-quadruplexes in RNA molecules alongside those in DNA. In contrast, RNase treatment significantly reduced the number of foci (35%), confirming that a substantial proportion of the detected G-quadruplexes are RNA-specific. These findings highlight the dual localization of G-quadruplexes in the genome and transcriptome of *H. volcanii*, emphasizing their potential roles in both nucleic acid stability and regulatory processes within these archaeal cells.

To broaden our approach to other archaea, we performed similar experiments using another archaea: *Thermococcus barophilus*, a hyperthermophilic archaea isolated from a deep-sea hydrothermal vent (70). Similar to *H. volcanii*, G4s were detected in both exponentially growing and stationary phase *T. barophilus* cells, with a significant increase in signal intensity in exponential phase cells (Supplemental Figure S3). Furthermore, treatment of *T. barophilus* cells with RNase A resulted in a significant decrease in G4 staining compared to untreated controls (Supplemental Figure S4).

### Modulation of G4s by a G4-ligand or helicases

We subsequently treated *H. volcanii* cells with PhenDC3, a highly specific G4-ligand already used in the *in vitro* assay. Cells were treated with PhenDC3 for two hours and subsequently processed for immunofluorescence analysis. We observed a statistically significant increase in G4 staining in PhenDC3 treated *vs*. untreated cells (Figure 5A).

**Fig. 5.**
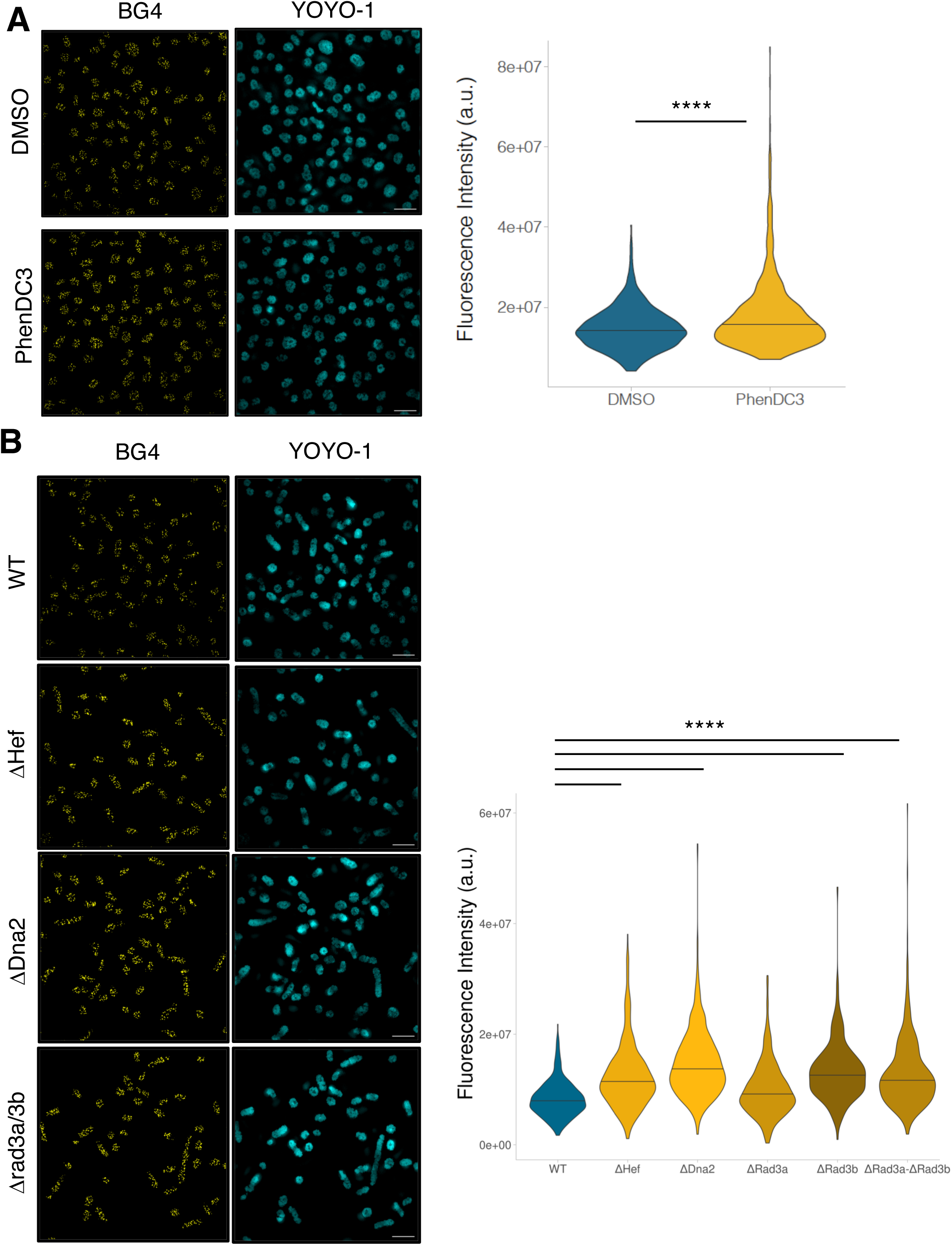
Chemical and genetic modulation of G4s in *H. volcanii*. (A) Quantification of total G4 fluorescence intensity in exponential phase *H. volcanii* cells treated with the G4 ligand, PhenDC3. The number of samples per condition is: DMSO=1578, PhenDC3=1159. (B). Quantification of total G4 fluorescence intensity in different helicase mutant *H. volcanii* strains. The number of samples per condition is: WT=449, ΔDna2=595, ΔHef=547, ΔRad3a=342, ΔRad3a-ΔRad3b=433, ΔRad3b=202. Statistical analyses were performed using a Mann-Whitney test. YOYO-1 was used to visualize nucleic acids. **** p > 0.0001. Bar, 5 µm.

To further characterize the presence and formation of G4s in our model archaeon, we aimed to identify potential helicases that may be involved in unwinding G4s in archaea. As such, we used different *H. volcanii* mutant strains that are deleted for genes encoding different helicases. Specifically, we focused on *H. volcanii* helicases whose eukaryotic putative orthologues had previously been proposed to regulate G4 dynamics *in vitro* and/or *in vivo*, such as Hef, Dna2 and Rad3a/3b (ref). These last two proteins share sequence homology in their helicase domain with SF2 helicases family members, including the G4-resolving helicases FANCJ (71), RTEL1 (72) and XPD (73), which are crucial for genome stability maintenance in higher eukaryotes. Genetic and imaging studies have revealed the central role of the archaeal helicase/nuclease Hef in restart of arrested replication forks (74–76). In mammals, the DNA2 helicase-nuclease has been proposed to play a crucial role in the resolution of G4s to prevent genome instability and replication stress (77). In addition to the *H. volcanii* Hef knock-out strain already available, we generated knock-out strains for Dna2 and Rad3a/3b to study their putative functions regarding G4 unwinding. Immunofluorescence experiments showed that there was a significant increase in DNA G4 staining in *H. volcanii* strains lacking these helicases compared to the wild-type strain (1.5-, 1.8-, and 1.5-fold for Δ*hef*, Δ*dna2,* and Δ*rad3a*/*rad3b*, respectively; Figure 5B). Only rad3a does not appear to be involved in G4 unwinding, suggesting that these two related helicases do not fulfill the same functions in the cell. We also analyzed the putative role of the ASH-Ski2 helicase in the G4 unwinding process in *T. Barophilus*. ASH-Ski2 is a Ski2-like helicase identified in archaea, characterized by conserved helicase motifs and is involved in RNA surveillance and decay pathways (78, 79). The absence of ASH-Ski2 expression led to a significant increase in the G4 signal, suggesting that this helicase may unwind G4 structures

## Discussion

Archaea, recognized as the third domain of life in the mid-1970s, have received relatively little attention in molecular and cellular biology compared to bacteria and eukaryotes. Yet, archaea are fascinating because their bacterial-like appearance contrasts with their enzymatic processes that more closely resemble those of eukaryotes (3). Despite the fact that many archaea thrive in extreme environments, they are also found in a wide range of “moderate” habitats, including the human microbiota (2) where their role in health and disease remains poorly understood and is currently being actively researched (80, 81). Recent analyses support a close relationship between eukaryotes and Asgard archaea, identifying them as the closest prokaryotic relatives of eukaryotes and offering key insights into the early stages of eukaryogenesis (6, 82). This underscores the importance of understanding archaeal biology more deeply, as it provides valuable insights into our evolutionary history.

Our study is the first to demonstrate the presence of G-quadruplexes (G4s), non-canonical nucleic acid structures (20, 36, 37), in archaea. G4s are among the most studied alternative nucleic acid structures of the last two decades. Several *in silico*, *in vitro*, and *in vivo* approaches have been developed to identify and validate G4-forming sequences across the genomes of various organisms; yet their role in evolution—and their own evolutionary dynamics—remain largely unexplored, although some pioneering studies have begun to address these questions (42, 83, 84). Our is the first to focus, in depth, on G4s in archaea, the closest living relatives of eukaryotes. Using publicly available annotated archaeal genomes and the G4Hunter algorithm, we were previously able to predict that G4s could be formed in the genome of every archaeal species examined (61). In this study, we focused primarily on *Haloferax volcanii*, a halophilic mesophilic archaeon (57). According to our G4Hunter algorithm, the *H. volcanii* genome contains approximately 1.55 putative G-quadruplex-forming sequences (PQS) per kilobase, which is of the same order of magnitude as the 1.2 PQS per kilobase observed in the human genome. Our biophysical experiments, conducted using various sequences from the *H. volcanii* genome with a G4Hunter score of at least 1.2, demonstrated that the majority of the tested sequences formed G4s under *in vitro* conditions. Importantly, these tested sequences were derived from not only from the *H. volcanii* main circular chromosome, but they were also taken from the two mega plasmids found within the strain used in our study (pHV1 and pHV3). This result suggests that G4s are found in all components of the *H. volcanii* genome, and are not limited to one specific element. Moreover, although we used classical ionic conditions corresponding to those found in human cells, it is highly likely that G4 structures can persist under the extreme intracellular ionic conditions characteristic of *H. volcanii*, even though only modest stabilization is typically observed with increased Na⁺ concentrations (unpublished observations).

To unequivocally demonstrate the presence of G4 structures in cells, we developed an immunofluorescence-based approach using the BG4 antibody and applied it to *H. volcanii* cells. Elegant super-resolution microscopy techniques have recently been used to visualize G4s and investigate their role in replication within mammalian cells (75, 76). Techniques like SIM and STORM overcome the diffraction limits of conventional fluorescence microscopy (56), offering nanometer-scale resolution and significant advantages for studying G-quadruplex structures in archaea. We were able to develop these protocols thanks to our prior work in detecting replication foci in *H. volcanii*, which provided a strong foundation for adapting super-resolution methods to archaea (52, 53).

Using SIM, we analyzed *H. volcanii* during exponential and stationary growth phases. G4s were clearly enriched in cells from the exponential phase. This likely reflects the increased DNA replication activity during exponential growth, where cells undergo continuous division. In contrast, stationary phase cells divide infrequently, explaining the reduced presence of G4s. We confirmed these results using the hyperthermophilic archaeon, *T. barophilus* (70). Nevertheless, it should be noted that archaea exhibit diverse ploidy levels, ranging from monoploidy to polyploidy, with some species maintaining multiple genome copies to enhance genetic stability and adaptability to extreme environments (85). The two models used in our study are polyploid and the greater number of G4s observed in the earlier growth phases could therefore be attributed to higher ploidy than in the stationary phase. One next step is to include a non-polyploid archaeon, such as *Sulfolobus solfataricus* (86), to compare the stability of G4 structures in different genomic contexts. Importantly, these results indicate that G4s can be formed in organisms that grow in considerably different conditions, implying an underlying prevalence of these structures not only archaea, but presumably throughout the three domains of life. Moreover, the detection of G-quadruplex structures in halophilic and thermophilic organisms suggest that they may contribute to molecular adaptation to extreme conditions of salinity or temperature. These approaches enable high-resolution detection of G4s, providing new insights into their roles in replication, transcription, and genome maintenance under extreme environmental conditions.

G4s were present at the DNA and RNA levels in both *H. volcanii* and *T. barophilus*, as indicated by a decreased G4 signal in cells following treatment with RNase A. This result is consistent with observations in eukaryotes, where G4s are known to also be formed not only in DNA, but also in RNA (87). In higher organisms, DNA G4s regulate a number of fundamental processes, such as DNA replication, transcription and telomere maintenance (16), whereas RNA G4s are thought to play a role in regulating translation and alternative splicing (29, 30). Interestingly, G4s are thought to regulate these processes either positively, through the recruitment of various initiating factors, or negatively, by acting as physical roadblocks preventing access to the appropriate molecular machinery (88). The presence of G4s in both RNA and DNA in archaea would suggest that these structures play similar functions in a diverse array of evolutionarily distinct organisms, underlying the fundamental nature and importance of G4s.Additionally, despite the presence of thousands of potential G4-forming sequences across the genome, only a limited number of BG4 foci are observed. This may indicate that the BG4 antibody preferentially detects clusters of G-quadruplexes (G4s) (89), rather than isolated structures—possibly due to higher local G4 density or enhanced stabilization within the cellular environment.

Initial detection of G-quadruplexes in archaea using immunofluorescence was a crucial first step, but genome-wide G4 mapping is now essential to comprehensively understand their distribution and potential regulatory roles in archaeal genomes. Beyond BG4-based approaches (90), several alternative strategies have recently emerged for the cellular investigation of G4 such as G4access (63), biomimetic ligands allowing selective G4 recognition and imaging in cells (91) and BrdU-tagged ligands (92). Some archaea also possess histones or histone-like proteins, suggesting a structured chromatin organization, albeit distinct from that of eukaryotes (11). Regarding G4 evolution, this raises the question of whether G4 structures are also located in open chromatin regions, as has been observed in mammalian cells (34, 63).

Having confirmed the presence of G4s in archaea, we aimed to identify potential players that are implicated in G4 stability/unfolding, using various helicase-deleted mutant strains of *H. volcanii* and one mutant in *T. barophilus*. Hef is a unique member of the XPF/MUS81 family of helicases/translocases and shares homologies with the human Fanconi Anemia protein FANCM, which has previously been implicated in destabilization of G4s, as human carcinoma cell lines lacking expression of FANCM were more sensitive to pyridostatin (PDS), a G4 ligand (74, 75, 93). Dna2 is a helicase/endonuclease that has been shown to bind and cleave G4s, primarily in telomeres both in yeast and human cell lines (77, 94). Rad3a and Rad3b are archaeal DNA helicases with strong sequence homology to eukaryotic XPD, sharing all the conserved helicase motifs and structural features characteristic of SF2 helicases that are also found in several eukaryotic G4-resolving helicases (FANCJ, DDX11, RTEL1) (71–73, 95). Interestingly, our current results have shown that loss of almost all of these helicases resulted in increased DNA G4s. This raises the intriguing possibility that there are multiple conserved helicases and enzymatic pathways that are implicated in regulating G4 stability and dynamics in archaea. Strikingly, the two Rad3, which have so far been little studied in archaea, do not appear to act in the same way on G4s. A similar finding has recently been reported for intermolecular gene conversion, suggesting very different roles for the two helicases (96).

Furthermore, it remains unclear whether these helicases cooperate in unwinding G4s. From a cell biology perspective, the involvement of multiple helicases in regulating G4 unfolding in DNA or RNA would be advantageous, as their cooperative action could compensate for the absence of one and prevent the accumulation of potentially deleterious G4s that threaten cell health and viability. Moreover, it has been observed in human cells that G4 DNA and RNA– DNA hybrids are both present within nucleosome-depleted regions, underscoring a pivotal role of G4 structures in R-loop formation (72, 97). G4 structures and R-loops are thought to contribute to genomic instability (98). These helicases could likely contribute to the resolution of G4-associated R-loops, thereby preserving genome stability and regulating gene expression in archaea similarly to their proposed role in eukaryotic systems. Future experiments investigating the colocalization of G4 structures using the BG4 antibody and R-loops would provide valuable insights into the spatial and functional relationship between these two structures in archaeal cells (99). In addition, the activities of these helicases towards G4 specifically could be analyzed using purified proteins (100). This would provide additional information on the roles of AshASH-Ski2 in *T. barophilus* (79) and Rad3a and Rad3b in *H. volcanii*.

The involvement of homologous human and archaeal helicases in G4 unfolding, along with the presence of G4s in both RNA and DNA in archaea, suggests an evolutionarily conserved mechanism for G4 formation and regulation. It also raises intriguing questions in evolutionary biology, indicating that archaea may possess molecular tools similar to those in eukaryotes for managing G4s. This is particularly notable given findings in *E. coli*, where RNA G4s are depleted, likely due to a lack of unfolding machinery. In fact, introducing RNA G4-forming sequences in *E. coli* impairs cell growth, implying an inability to resolve these structures (69). However, the role of G4s in bacteria could be much more complex, as they may be involved in regulating translation and may be effective targets for G4 ligands in both *E. coli* and *H. pylori*. (51, 101, 102)

It is now widely accepted that G4s can play antagonistic roles: promoting some molecular processes, but also interfering with them. The origin of this duality can be investigated by studying prokaryotic organisms. Until now, there has been virtually no data on the detection and function of G4s in archaea. This work opens up new perspectives for understanding the relevance of G4s and how they influence molecular processes such as replication, transcription, translation, and repair. The identification of potential helicases involved in G4 unwinding provides a tool for studying their impact, although the significance of these proteins *in vivo* remains to be better defined.

Overall, our work is the first unequivocal experimental demonstration of G4 formation in archaea, an exciting result that opens the door to further studies that will help to answer fundamental questions about the evolution of organisms and the molecular mechanisms regulating DNA structure, dynamics, and the regulation of gene expression. To fully understand the diversity and conservation of G4 structures in archaea, it is now essential to significantly expand the repertoire of archaeal species under investigation—particularly those most closely related to eukaryotes.

## Acknowledgements

We thank M. Imezar, H. Becker, L. Mellottee, T. Garnier, J. Gros, S. Duigou (LOB), JC. Andrau (IGMM) and L. Lacroix (ENS) for technical assistance and helpful discussions.

## Funding

ANR G4Access [ANR-20-CE12-0023]

ANR Morphoscope2 [ANR-11-EQPX-0029] ANR France BioImaging [ANR-10-INBS-04]

## Data and materials availability

All data needed to evaluate the conclusions in the paper are present in the paper and/or the Supplementary Materials.

## Author contributions

Conceptualization: ZA, JLM, LG

Methodology: ZA, KS, VB, RL, NO, BCO, DV, TA, PM, JLM, LG

Investigation: ZA, AC, GV, DN, RZ, TJ, NV, OP, KS, VB, MK, MB

Supervision: JLM, NO, RL, LG

Writing—original draft: JLM, ZA, LG

Writing—review & editing: JLM, ZA, LG

