## Supplemental Figures for "Archaeal G-Quadruplexes: A Novel Model for Understanding Unusual DNA/RNA Structures Across the Tree of Life"

### Supplemental Figure 1

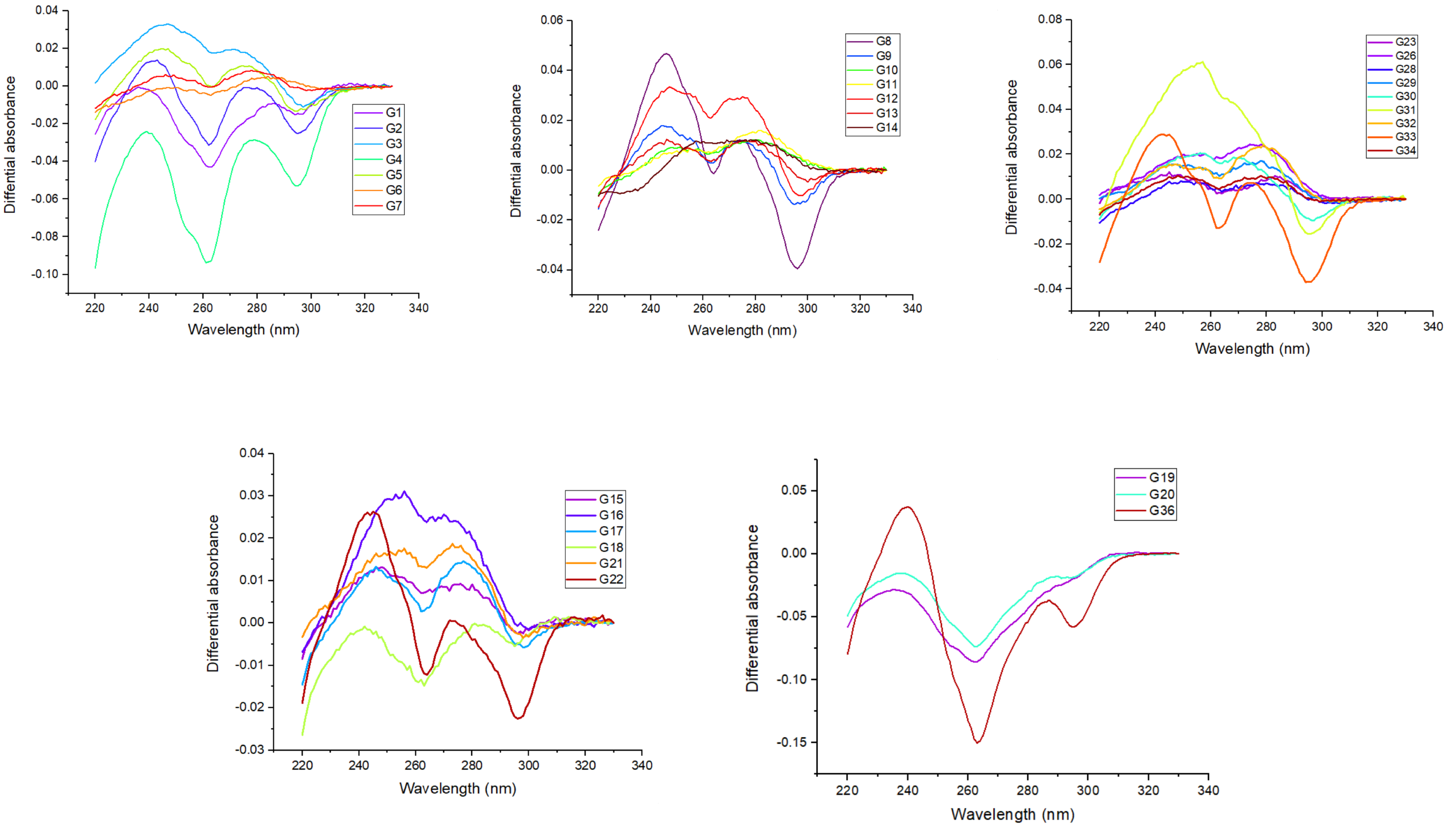

### Supplemental Figure 2

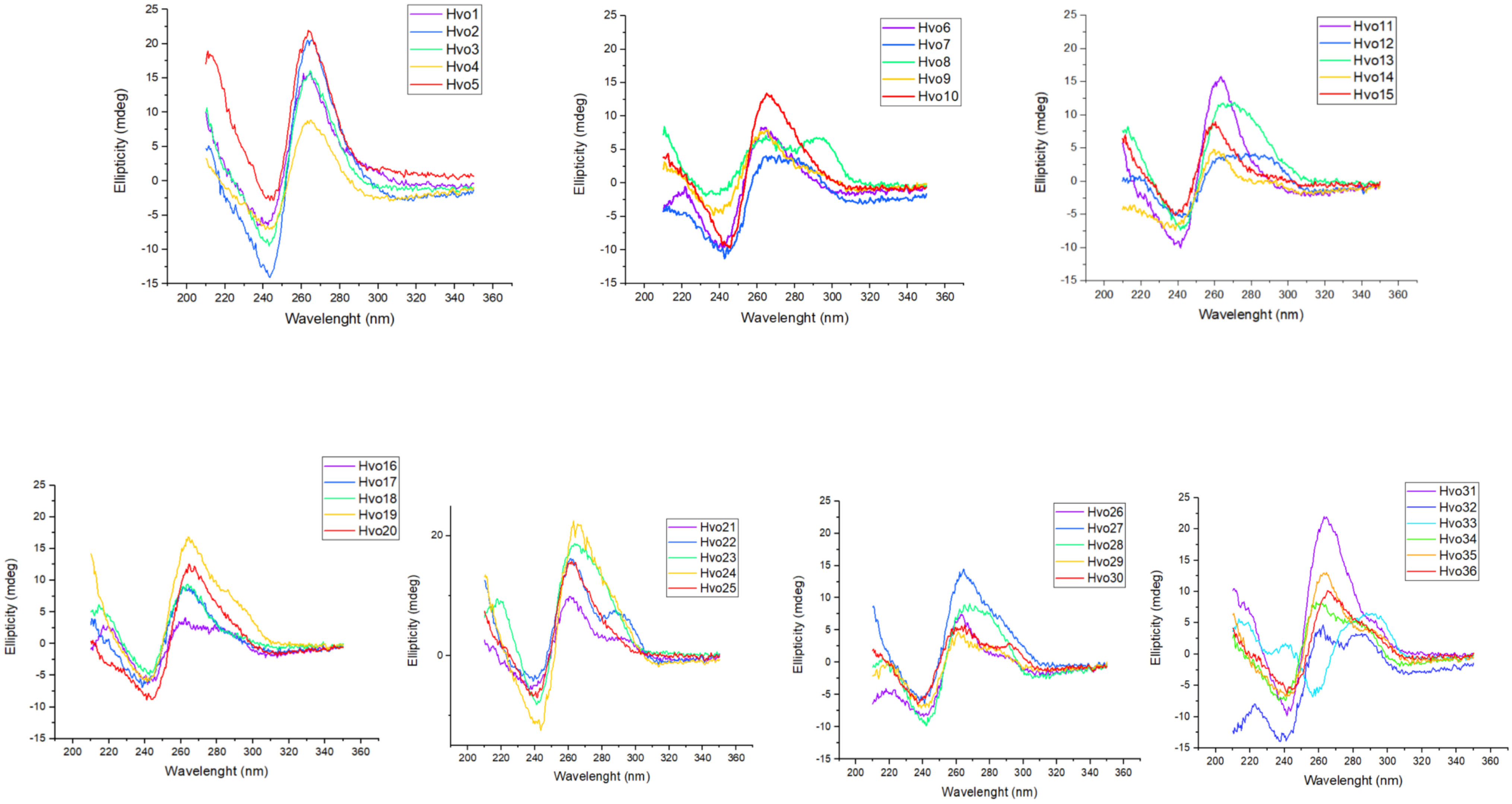

**Supplemental Figure 3**

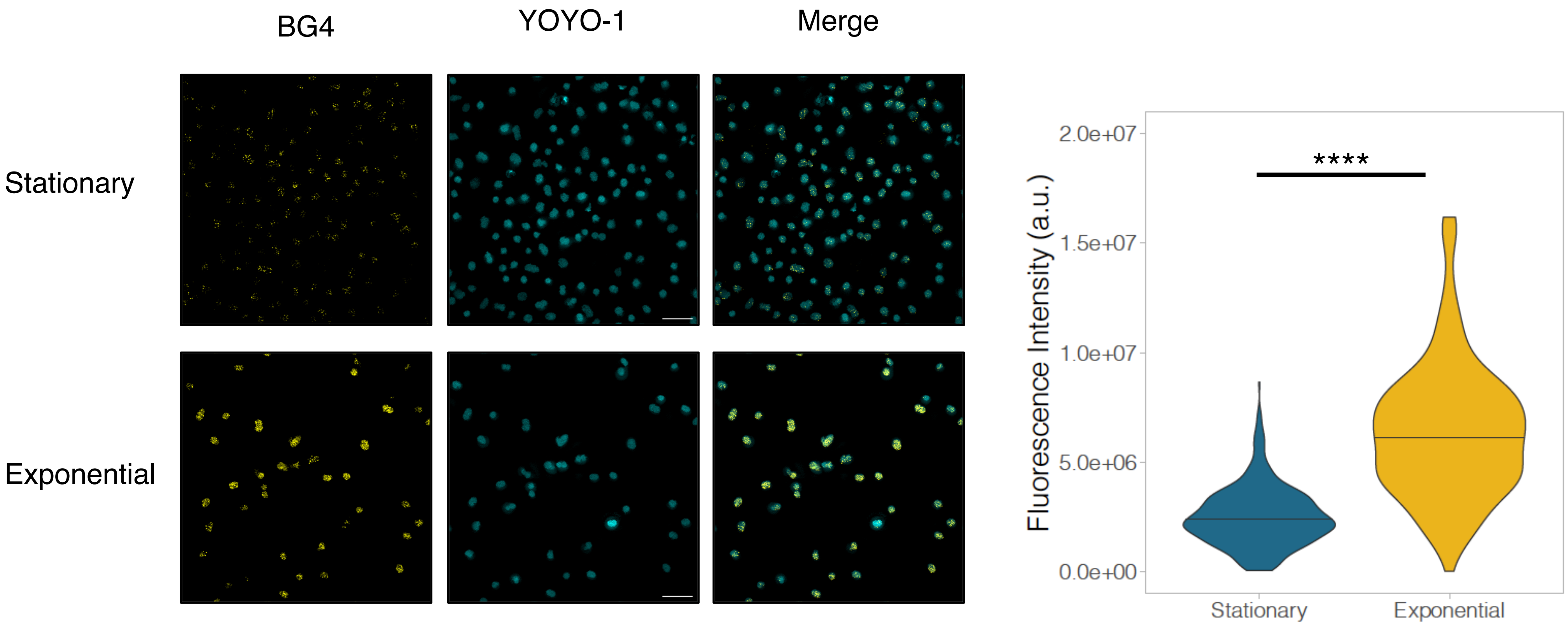

### Supplemental Figure 4

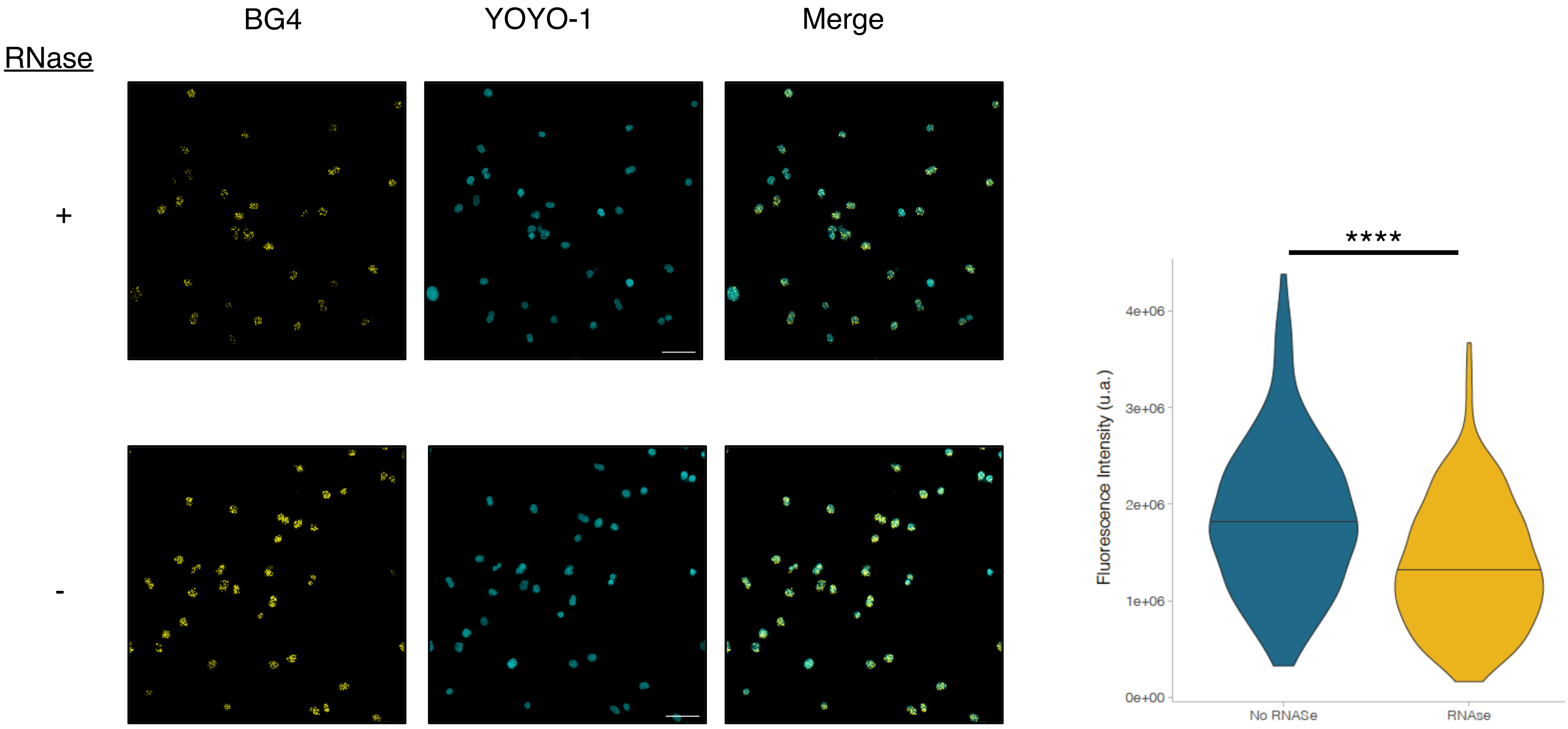

**Supplemental Figure 5**

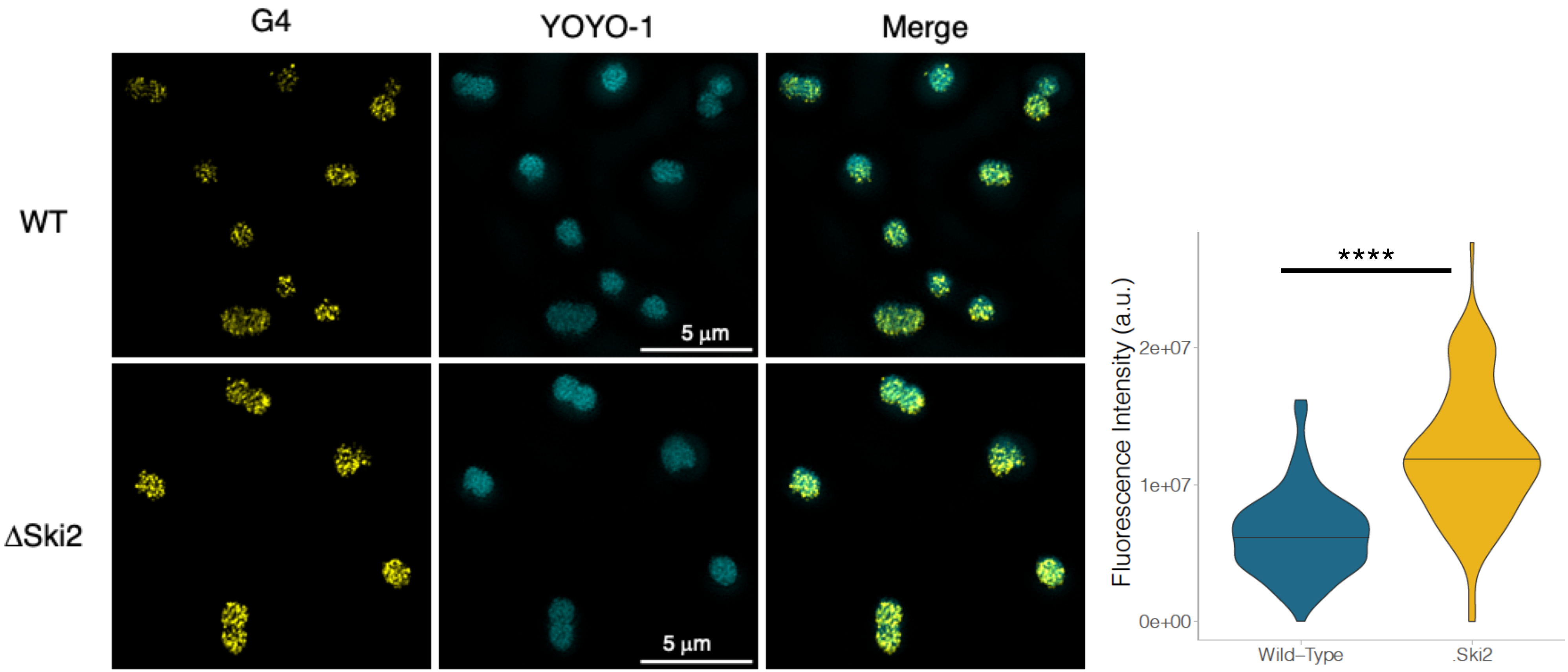
