## Supplemental Tables and Legends for "Archaeal G-Quadruplexes: A Novel Model for Understanding Unusual DNA/RNA Structures Across the Tree of Life"

**Supplemental Table 1.** DNA oligonucleotides from *H. volcanii* used to study G4 folding. All sequences are provided in the 5' → 3'

| Name | Sequence | G4H score | Localisation |
|---|---|---|---|
| Hvo-G1 | CGGGGTCGGGACTCGGCATCGTGGTGGGGT | 1,61 | chr |
| Hvo-G2 | GGGGTGGATGGGCGACGCGACGAAGGGG | 1,61 | chr |
| Hvo-G3 | TGGGGGTGAGGATACAGACGGCATCGGGGGT | 1,59 | chr |
| Hvo-G4 | GGGGTGACCGAACCTAATGGGGAAGGGGGGAAGGGGGGAT | 1,46 | chr |
| Hvo-G5 | CGGGGGAAAAAGGTTCAGCGGTCGGGGAG | 1,45 | chr |
| Hvo-G6 | CGGGCACCGGGACTCACGGAGGGGAGGGC | 1,44 | chr |
| Hvo-G7 | CGGGGTGTGGGCACGGTCGGCAGACGGGC | 1,44 | chr |
| Hvo-G8 | CGGGTTGGTCGGGGAGTCCGGTTGCGGGA | 1,41 | chr |
| Hvo-G9 | CGGGGGTCGGGCATCGGTGTTCATCGGGC | 1,41 | chr |
| Hvo-G10 | CGGGGAGCGACGGCGACGGCGACGGGGA | 1,41 | chr |
| Hvo-G11 | GACGGGGCGGGGTCACTCCGGCCGGGGGTC | 1,40 | chr |
| Hvo-G12 | CGGGAAGGGTTGACGGTACGCTCGGGGC | 1,38 | chr |
| Hvo-G13 | GGGGTGGGCGAGGAACTCGACGCGGG | 1,38 | chr |
| Hvo-G14 | CGGGGGAGTCCGGCGCGGGCGCTCGGGA | 1,38 | chr |
| Hvo-G15 | AGGGGATGGATCGGCAGGAGGACGGT | 1,25 | pHV1 |
| Hvo-G16 | AGGAATCGGGAATGGACTGGGGCCGGA | 1,24 | pHV1 |
| Hvo-G17 | GTCGGTGGCGGGACCGGGAAGGGTA | 1,20 | pHV1 |
| Hvo-G18 | CGGTGGCGGGACCGGGAAGGGTAAGA | 1,15 | pHV1 |
| Hvo-G19 | AGGGCTGCGGACAGGGCGGGATGATGGGT | 1,41 | pHV4 |
| Hvo-G20 | CGGGCATCGTCGGGACAGGTGGGCGGGC | 1,38 | pHV4 |
| Hvo-G21 | CGAGGTCGGGGCGGTCGGTCGCGGGGATGGTCA | 1,38 | chr |
| Hvo-G22 | GGGTGGCGGCGGCGGTGGCGGCATGGG | 1,37 | chr |
| Hvo-G23 | CCAGGGTAGGAAGACGGACCACGGGGAGGGA | 1,37 | chr |
| Hvo-G24 | AGGGGCTATCGGTGGCTGTCGCGGGGT | 1,37 | chr |
| Hvo-G25 | TGGGGGTCCTGGTCGGGGCCGACCGGGAT | 1,37 | chr |
| Hvo-G26 | TGGGCCATCGGCGCGGGGTCGTCGGGGT | 1,36 | chr |
| Hvo-G27 | CCGTGGGGGTCCTCGGGATTGTAGGGTGCGGGC | 1,35 | chr |
| Hvo-G28 | CAGGAGTCGGAGGACTACGAGGGCGGGGCC | 1,35 | chr |
| Hvo-G29 | CGGGCAACGGTCGAGGTGGTGACGGGGAC | 1,35 | chr |
| Hvo-G30 | AGGGGTCGGCAGGGTCCGGTATCAGGGTA | 1,35 | chr |
| Hvo-G31 | TGGGAAGGATGGGCCGAGGACGAGGGGAT | 1,34 | chr |
| Hvo-G32 | CGGGTGACGACGGGCCGGCGGTCGGGGAC | 1,33 | chr |
| Hvo-G33 | TGGGAGAAGGGCGCGGGCAAGATTTGGGC | 1,33 | chr |
| Hvo-G34 | AGGGGACCGGGTGGTCGCCGGGGAGTC | 1,31 | chr |
| Hvo-G35 | TGGGTCGGGGCGAGCGTTCGGGGGCGGT | 1,32 | chr |
| Hvo-G36 | CGGGCATCGTCGGGACAGGTGGGCGGGC | 1,38 | pHV4 |

**Supplemental Table 2**. DNA oligonucleotides, positive and negative controls for G4 folding, used in the present study. All sequences are provided in the 5' → 3'

| Name | Sequence | Length | Topology |
|---|---|---|---|
| c-kit* | GGCGAGGAGGGGCGTGGCCGGC | 23 | Anti-parallel |
| Bcl2 | GGGCGCGGGAGGAATTGGGCGGG | 23 | Hybrid |
| 23TAG | TAGGGTTAGGGTTAGGGTTAGGG | 23 | Hybrid |
| ckit2 | CGGGCGGGCGCTAGGGAGGGT | 21 | parallel |
| 26CEB | AAGGGTGGGTGTAAGTGTGGGTGGGT | 26 | Parallel |
| 22CTA | AGGGCTAGGGCTAGGGCTAGGG | 22 | Anti-Parallel |
| Ds-lac | GAATTGTGAGCGCTCACAATTC | 22 | Hairpin |
| Hp2 | TCGGTATTGTGTTTCACAATACCGA | 25 | Hairpin |
| F21T (double labeled) | GGGTTAGGGTTAGGGTTAGGG | 21 | Hybrid |

**Supplemental Table 3.** Oligonucleotides used in the present study to generate Rad3a/b knock-out strains by the SLIC method and the Dna2 knock-out strain. All sequences are provided in the 5' → 3' (US : Upstream and DS : Downstream)

| Name | Sequence |
|---|---|
| US-F rad3b | TCGACGGTATCGATAAGCTTGATATCGAATTGACGACCCAGAGGACGACGCC |
| US-R rad3b | GGGAGAATCTCGGCCGCCATC |
| DS-F rad3b | TTTTAATCGGATGGCGGCCGAGATTCTCCCGCCTGAGCGGGTCGCGCTCGG |
| DS-R rad3b | AAAAGCTGGAGCTCCACCGCGGTGGCGGCCTCATCATGCCGGCCATCGTCG |
| US-F rad3a | TCGACGGTATCGATAAGCTTGATATCGAATTTTCACTGGCTCGGCGACGACGA |
| US-R rad3a | AGGGGTGTCGGTGACTGGTCGCGGA |
| DS-F rad3a | TACGGGTCCGCGACCAGTCACCGACACCCCTTCGCGGCCGGGGTCGCGCCG |
| DS-R rad3a | AAAAGCTGGAGCTCCACCGCGGTGGCGGCCCGACGCGCAGGTGTGGATTCCACAG |
| dna5f | GCCTATCGTCTAGATCCGTGTATTGG |
| dna5r | GCGAGGATCCCTTTCTCGCCCACCCG |
| dna3f | GACCGGATCCTCGCGTGGGCGCGGCG |
| dna3r | CGTCGTGGGTACCGAACTCGACCGCG |

**Supplemental Table S4**

| Name | FRET MC | ThT | NMM | IDS | CD | Conclusion |
|---|---|---|---|---|---|---|
| Hvo-G1 | | | | | | |
| Hvo-G2 | | | | | | |
| Hvo-G3 | | | | | | |
| Hvo-G4 | | | | | | |
| Hvo-G5 | | | | | | |
| Hvo-G6 | | | | | | |
| Hvo-G7 | | | | | | |
| *Hvo-G8* | | | | | | |
| *Hvo-G9* | | | | | | |
| Hvo-G10 | | | | | | |
| Hvo-G11 | | | | | | |
| Hvo-G12 | | | | | | |
| *Hvo-G13* | | | | | | |
| Hvo-G14 | | | | | | |
| *Hvo-G15* | | | | | | |
| *Hvo-G16* | | | | | | |
| *Hvo-G17* | | | | | | |
| *Hvo-G18* | | | | | | |
| *Hvo-G19* | | | | | | |
| *Hvo-G20* | | | | | | |
| *Hvo-G21* | | | | | | |
| *Hvo-G22* | | | | | | |
| *Hvo-G23* | | | | | | |
| *Hvo-G24* | | | | | | |
| *Hvo-G25* | | | | | | |
| *Hvo-G26* | | | | | | |
| *Hvo-G27* | | | | | | |
| *Hvo-G28* | | | | | | |
| *Hvo-G29* | | | | | | |
| *Hvo-G30* | | | | | | |
| *Hvo-G31* | | | | | | |
| *Hvo-G32* | | | | | | |
| *Hvo-G33* | | | | | | |
| *Hvo-G34* | | | | | | |
| *Hvo-G35* | | | | | | |
| *Hvo-G36* | | | | | | |

| | |
|---|---|
| | G4 forming |
| | Likely to form G4 (unstable or at low propensity) |
| | Unlikely to form G4 |
| | Non determined |

**Supplemental figure legends**

**Supplemental Fig. 1.** Isothermal difference spectra (IDS) of the *H. volcanii* sequences.

**Supplemental Fig. 2.** Circular dichroism spectra of the *H. volcanii* sequences.

**Supplemental Fig. 3.** Immunofluorescence labelling and quantification of G4s in fixed *T. barophilus* cells in exponential and stationary phase. Statistical analyses were performed using a Mann-Whitney test. **** $p > 0.0001$. Bar, 5 μm.

**Supplemental Fig. 4.** Immunofluorescence labelling and quantification of G4s in fixed *T. barophilus* cells in exponential phase treated with RNase A. Statistical analyses were performed using a Mann-Whitney test. **** $p > 0.0001$. Bar, 5 μm.

**Supplemental Fig. 5.** Immunofluorescence labelling of G4s in WT or ΔSki2 *T. barophilus* exponential phase cells. Bar, 5 μm.
